# The kinectome: a comprehensive kinematic map of human motion in health and disease

**DOI:** 10.1101/2021.10.05.463174

**Authors:** Emahnuel Troisi Lopez, Pierpaolo Sorrentino, Marianna Liparoti, Roberta Minino, Anna Carotenuto, Enrico Amico, Giuseppe Sorrentino

## Abstract

Effective human movement requires the coordinated participation of the whole musculoskeletal system. Here we propose to represent the human body movements as a network (that we named “kinectome”), where nodes are body parts, and edges are defined as the correlations of the accelerations between each pair of body parts during gait. We apply this framework in healthy individuals and patients with Parkinson’s disease (PD). The network dynamics in Parkinson’s display high variability, as conveyed by the high variance and the modular structure in the patients’ kinectomes. Furthermore, our analysis identified a set of anatomical elements that are specifically related to the balance impairment in PD. Furthermore, each participant could be identified based on its kinectome patterns, akin to a “fingerprint” of movement, confirming that our approach captures relevant features of gait. We hope that applying network approaches to human kinematics yields new insights to characterize human movement.

## Introduction

Human voluntary movement requires spatially and temporally coordinated activations of specific muscular groups (*1, 2*), resulting in complex patterns, whose description requires taking into account a large number of interactions concurrently (*3*). Such patterns are fine-tuned, and small changes may lead to significant effects (*4*). As an example, the upright posture requires extremely refined control mechanisms in order to keep balance during gait (*5, 6*). Among such mechanisms, the different musculoskeletal segments developed hierarchical relationships (*7*). Therefore, an accurate characterization of movement requires precise measurements and appropriate mathematical methods, in order to capture small but physiologically relevant changes in movement patterns.

To date, human movement, and notably gait, has been thoroughly studied focusing on specific body elements, or summarising multiple synergies into few synthetic parameters (*8*–*11*), that often describe the overall movement rather than the complex patterns of interactions that generated it. Indeed, since the human body is highly connected, whole-body interactions are needed to provide a comprehensive account of the complex movement dynamics (*12, 13*), on top of the fixed musculoskeletal structure. To overcome this limitation, network science, including graph theory (*14*), may provide useful tools to describe such complex patterns. Indeed, in the last decades, network science has been a major tool in exploring complex systems, and has allowed us to effectively represent them (*15*). Using the network science viewpoint, we can represent anatomical elements as nodes, and their relationships throughout movements as edges, defining the “network of human movement”.

Recently, the first applications of network analysis to the study of the human body movement proved successful. Utilizing electromyography, Boonstra et al analysed the leg muscles network, highlighting the presence of lower and higher frequency components, related to between and within legs connectivity, respectively (*16*). The authors suggested that network analysis may be suitable to study the motor system also in clinical conditions. Moreover, a combined musculoskeletal network structure was investigated by Kerkman et al., which examined in depth the different frequency-specific networks during postural control (*17*). The study showed that the examined network presented frequency-specific relationships with the synaptic input to motor neurons. Yet, despite these first efforts in bridging the gap between network and motor sciences, to date there is no comprehensive description of the kinematics of movement.

Here, we set out to identify the large-scale characteristics of the human gait. Indeed, human locomotion requires the coordination of the whole body, and cannot be accounted for by the lower limbs alone (*18, 19*). Hence, we considered the whole body as an integrated and synergistic system, whose individual elements are in a constant and reciprocal biomechanical relationship, constrained by the individual anatomical characteristics. To this end, we performed a three-dimensional motion analysis using a stereophotogrammetric system, which is the gold standard for quantitative analysis of movement (*20*), and widely applied for the assessment of motor skills in health and disease (*10, 21*–*24*). Specifically, we captured the position of reflective markers applied on specific bone reference points during gait. Each bone marker was considered as a node, and the edges linking these nodes were defined by the covariance of the acceleration and jerk (i.e., the first derivative of acceleration with respect to time) between each pair of bone markers. We name the resulting network the human “*kinectome*”.

In this work, we explore the features of the human kinectome in a cohort of healthy subjects (HS). Furthermore, in order to explore the applicability of our framework to a clinical setup, we compared the kinectomes of individuals affected by Parkinson’s disease (PD), a neurodegenerative disorder which disrupts the motor patterns of the patient (*25*), to those of a group of matched healthy controls (HC). We hypothesized that PD patients would be less capable of maintaining the (presumably) optimal motor strategy seen in the HC. According to this hypothesis, we first explored the structure of the kinectomes, expecting a dysregulated (i.e., more variable) organisation in patients compared with controls. Secondly, to test the reliability of the kinectome in health and disease, we performed an identifiability (*26*) analysis, in order to check whether we could identify subjects by relying on their kinectomes, similarly to a “motion fingerprint”. This idea came from recent evidence that a dysregulated activity would make identifiability harder in patients (*27*). Finally, to check the clinical validity of our framework, we related nodal topological features of the kinectome to the clinical disability, measured using the Unified Parkinson’s Disease Rating Scale part III (UPDRS) (*28*), and with the stability of gait, measured with the Trunk Displacement Index (TDI) (*29*).

## Results

We investigated the covariance matrices of the motor patterns adopted during gait, i.e., the *kinectomes* (Fig. 1), constructed from the acceleration and jerk time series in the mediolateral and anteroposterior axes, in three groups: healthy individuals (HS), and subjects affected by PD (PD) and matched healthy controls (HC). Kinematic time series were collected from bone markers applied on the skin of the participants (Fig. 1A), through a stereophotogrammetric system. We aimed to test the hypothesis that kinectomes could serve as a global descriptor of human gait kinematics, both in health and disease.

**Figure 1.**
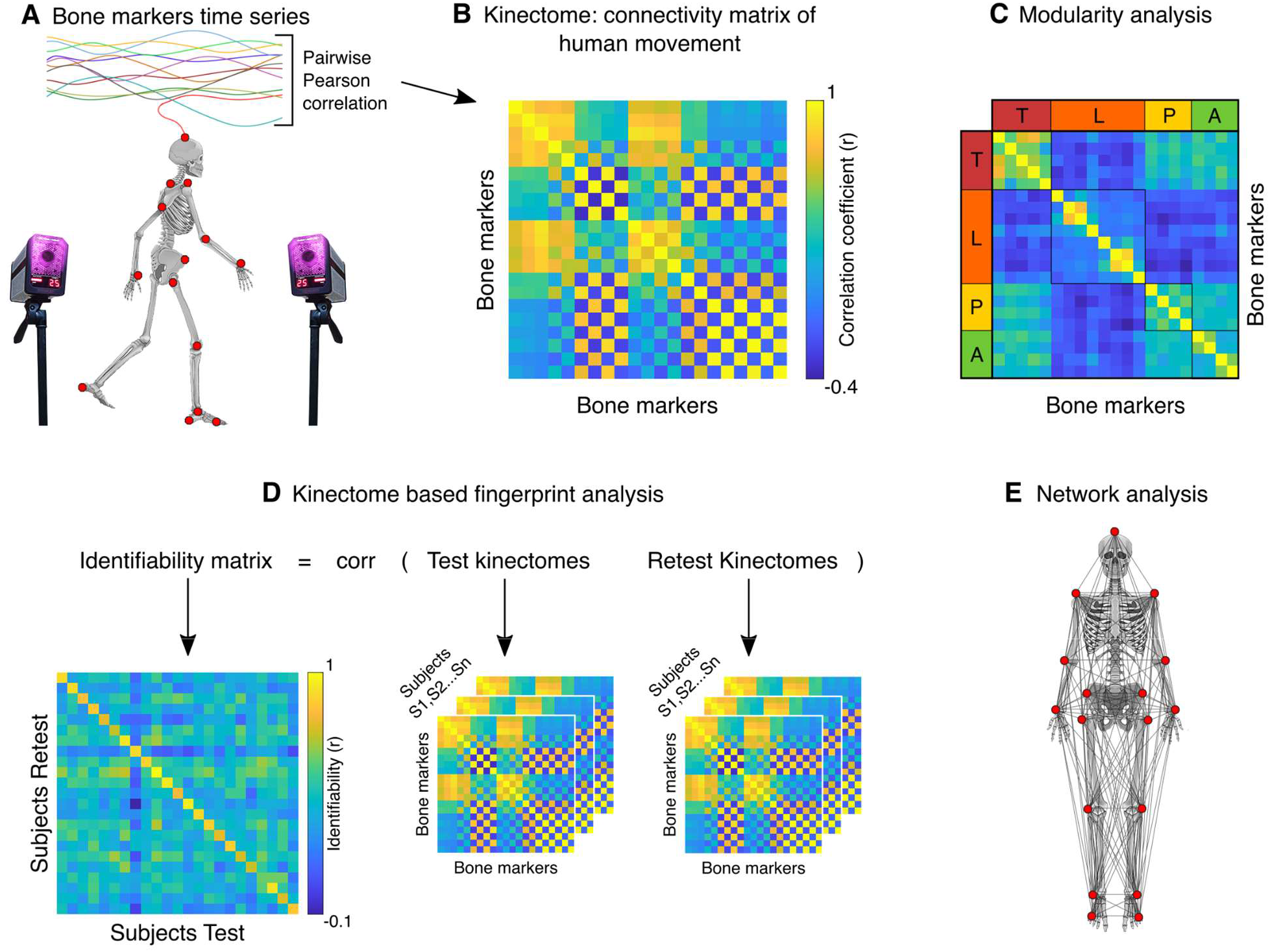
Scheme of kinectome analysis. **(A)** Markers positions on the bone landmarks. Acceleration and jerk time series are computed based on the positions of the markers during the gait cycle, recorded by a stereophotogrammetric system. **(B)** Kinectomes: the covariance matrix computed correlating each pair of the bone markers time series; different kinectomes were built, based on the mediolateral and anteroposterior axis, separately taking into account the accelerations and the jerks during gait. **(C)** The functional network modularity was investigated through the Louvain method, an algorithm employed for community detection. **(D)** Schematic illustration of the fingerprint analysis. Two kinectomes (named test and retest) have been computed for each subject. The Identifiability matrix is obtained correlating the test and retest kinectomes of each subject. The main diagonal displays self-identifiability. **(E)** Graphical representation of the bone markers network used for the topological analysis. Note that the bone markers positioned on the back of the body are not visible.

### Kinectomes characteristics

We started from a group-level analysis comparing the average kinectomes of healthy subjects and PD patients. Specifically, after building subject-specific kinectomes (Fig. 2A), we averaged them within each group, obtaining the group-specific (i.e., HS, HC, PD) kinectomes in which each element was the group-averaged correlations between the time-series derived from two bone markers during gait. Then, we compared the average values of the kinectomes in HC and PD patients. The analysis was performed for both the anteroposterior and mediolateral axes, and for both acceleration and jerk-based time series (Fig. 2B). However, neither acceleration nor jerk kinectomes highlighted any significant difference between the two groups. That is, the acceleration and jerk patterns of the two groups were similar to each other.

**Figure 2.**
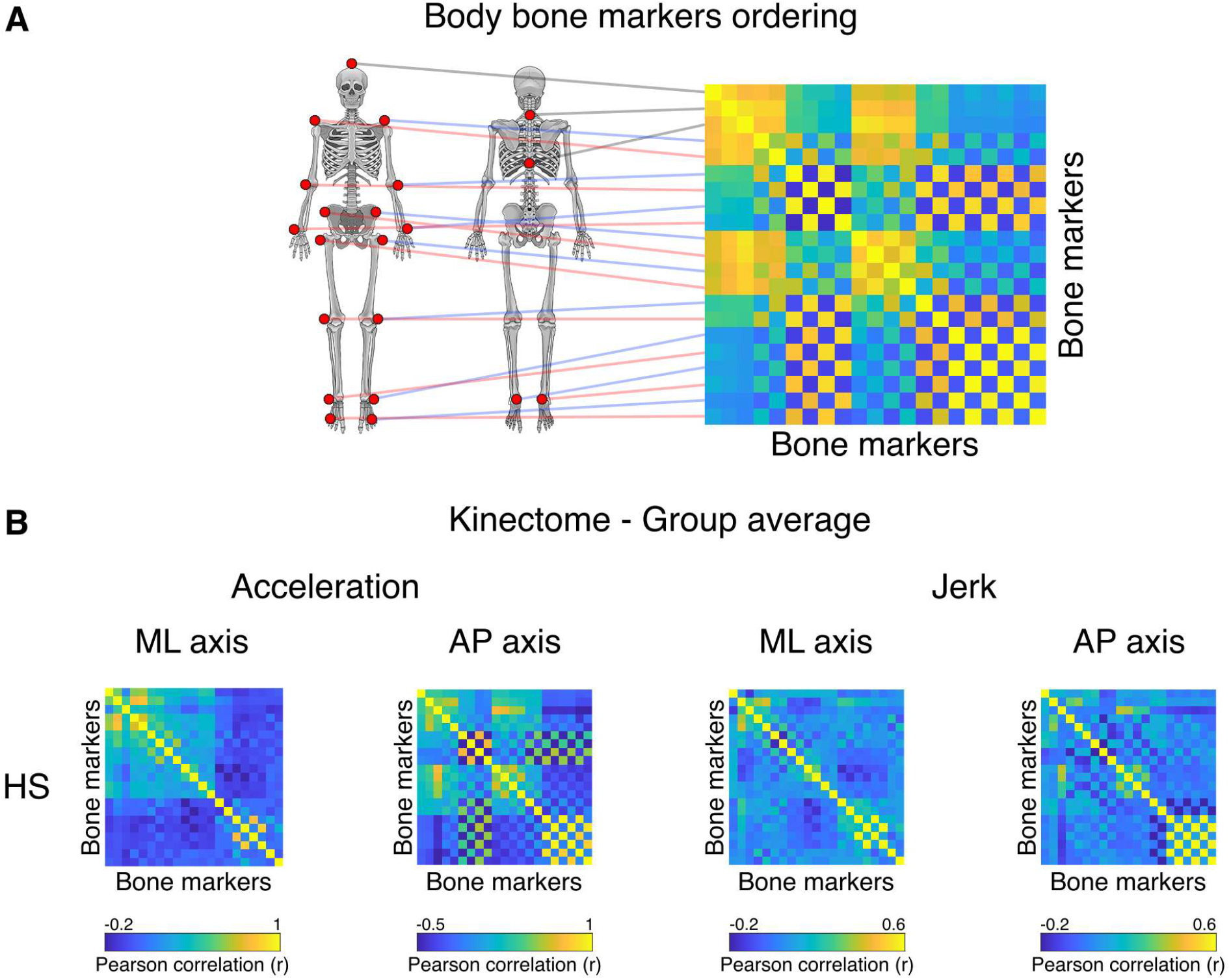
From bones to kinectomes. **(A)** Illustration of the bone markers position on the kinectome. Each bone marker kinematic information is used as entry data for both rows and columns. The elements of the kinectome stem from the pairwise interaction between bone markers. **(B)** Acceleration and jerk kinectomes averaged among healthy subjects (HS) in the mediolateral (ML) and the anteroposterior (AP) axes. The interactions between body elements varies according to both the specific axis and measurement (acceleration or jerk) taken into account.

Next, we checked the kinectomes’ within-group variability, by evaluating the standard deviation of kinectomes across HC and PD patients (Fig. 3). Notably, the variability in the whole-body movement patterns between the two groups showed higher standard deviation among PD patients in the anteroposterior acceleration (p = 0.0002, Bonferroni cut-off p < 0.0125), compared to the HC group. Figure 3 shows the considerable variability expressed by the PD patients in the anteroposterior acceleration. This suggests augmented variability in movement patterns belonging to the PD population.

**Figure 3.**
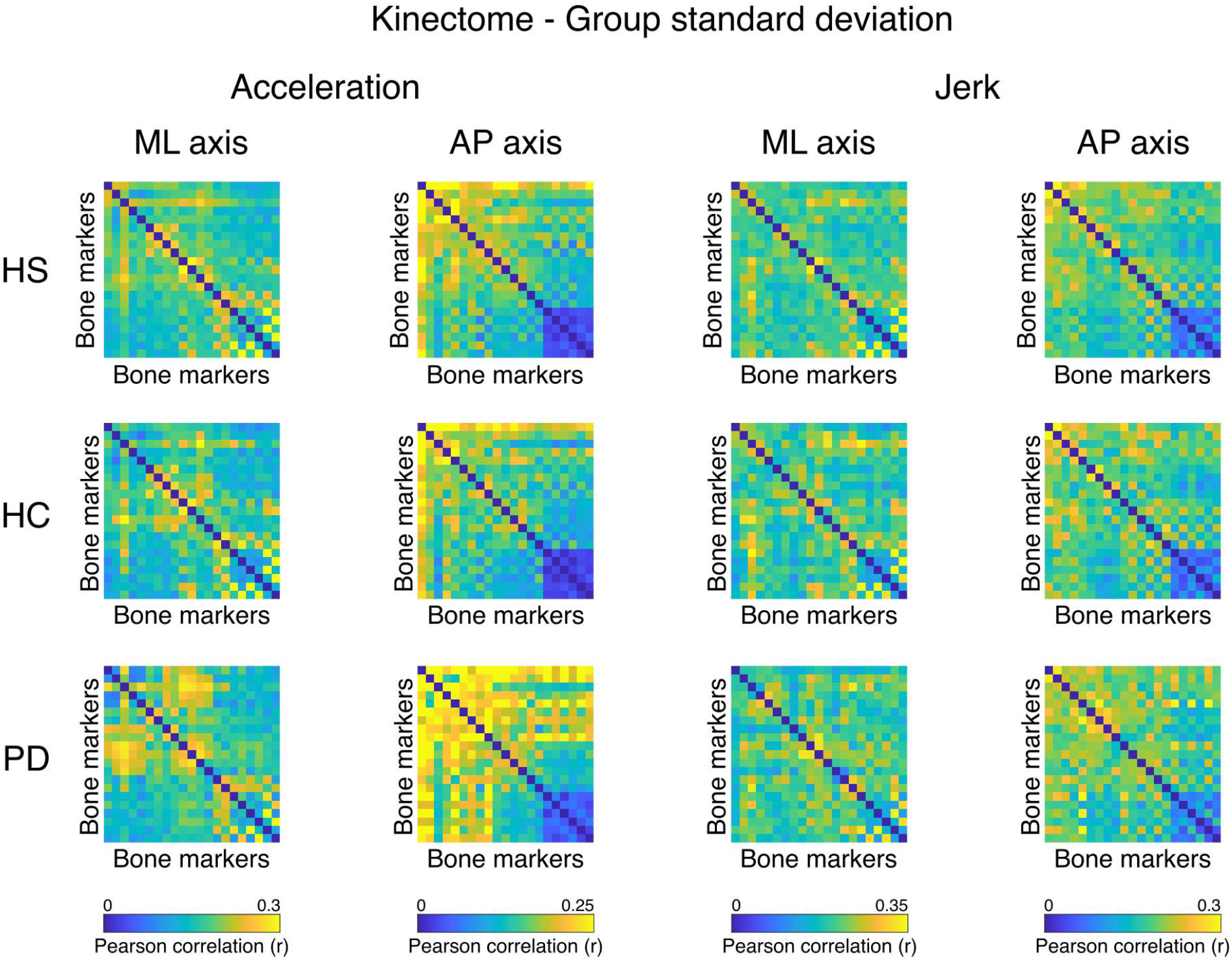
Within group heterogeneity of the kinectomes. Standard deviations of the acceleration and jerk kinectomes of the three groups (healthy subjects (HS), healthy controls (HC), Parkinson’s patients (PD). Both mediolateral (ML) and anteroposterior (AP) axes are shown. Higher values (i.e., yellow entries in the matrices) indicate greater heterogeneity.

### Modularity analysis

We then set out to provide a principled description of the kinectomes’ topological structure, by investigating the emerging modular structure of the kinectomes (see Methods for further details). To this end, we computed the allegiance matrices (*30*), which contain the probability of any two bone markers being clustered in the same community across individuals. This means that two or more bone markers belonging to the same cluster refer to body parts which are likely to coordinate themselves toward the same motor pattern. Figure 4 shows that the HS and HC groups share the same communities, while the PD group features a different clustering pattern. Specifically, in the healthy groups, the ML allegiance matrix showed three communities: *1)* upper trunk and arms, *2)* head, forearms and pelvis, *3)* legs and feet; in the PD group, the same matrix showed four different communities: *1)* upper trunk and right upper arm, *2)* head, left upper arm, forearms and pelvis (upper portion), *3)* legs and feet, *4)* pelvis (lower portion). The AP allegiance matrix in the healthy groups highlighted three communities: 1) head, upper trunk and pelvis, 2) left leg and foot, and right upper arm and forearm, *3)* right leg and foot, and left upper arm and forearm; in the PD group, the same matrix identified four communities: *1)* head and upper trunk (upper portion), *2)* left leg and foot, and right upper arm, *3)* upper trunk (lower portion), pelvis and right forearm, *4)* right leg and left upper arm and forearm.

**Figure 4.**
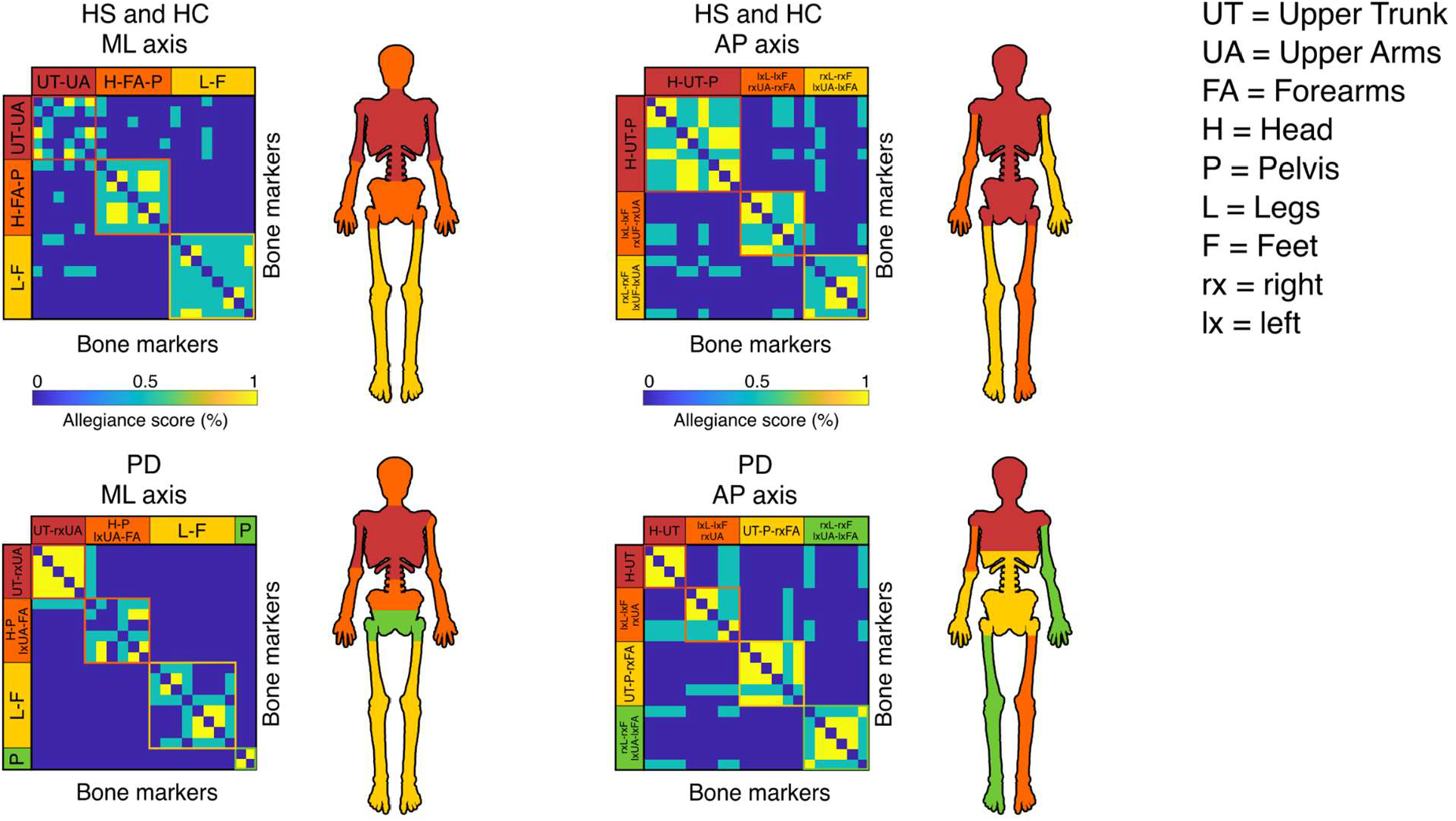
Kinematic modular organization of the kinectomes. Allegiance matrices for cluster analysis, based on the Louvain method and stabilised through 100 iterations. The algorithm automatically defines which body parts belong to the same community, suggesting a functional relationship among those elements. Each matrix includes clustering information from both accelerations and jerks. Healthy subjects (HS) and healthy controls (HC) share the same communities in both mediolateral and anteroposterior axes. Parkinson’s disease (PD) patients’ matrices show a different structural organisation. Body parts depicted with the same colour belong to the same functional community.

This approach allowed us to observe how the body kinematic is organised during gait, both in health and disease. It is noteworthy that the algorithm calculating the communities split the body parts symmetrically in healthy individuals, while the same result was not achieved for the PD patients, which might be related with the typically asymmetrical motor impairment in PD.

### Fingerprint of human movement

Can we identify humans based on their motion patterns? In other words, is there a “motion fingerprint” inherent to the kinectomes? To answer this question, we tried to identify individuals through their kinectomes, obtained from different gait sessions recorded during the same day. To this aim, we started by building an identifiability matrix based on kinectomes (*26*). In the identifiability matrix, the rows refer to kinectomes of the first recording (Test kinectomes in Fig. 1D), and the entries on the columns refer to the kinectomes of the second recording (Retest kinectomes in Fig. 1D). The entries of the Identifiability matrix are Pearson’s correlation coefficients between the kinectomes derived from the first and the second recording sessions. Briefly, from the Identifiability matrix it was possible to calculate three parameters: I-self (self-similarity across the two sessions), I-others (similarity with other individuals within the group), and I-diff (differential identifiability), obtained subtracting the I-other from the I-self. Figure 5A displays the main results of the fingerprint analysis, focusing on the jerk-based identifiability matrix (see Fig. S2 for acceleration kinectomes identifiability). The identifiability matrix was firstly evaluated in the healthy population; thereafter, the same matrix was assessed in PD and matched control populations. AP and ML jerk resulted to be the best quantities for the gait identifiability (highest I-diff values) and this result is stable across the three groups. Specifically, the ML Jerk in the HS group showed an I-diff equal to 37.8% (p < 0.0001), with a success rate (SR) to identify the subjects equal to 99.87%; AP jerk showed an I-diff and a SR rate equivalent to 25.89% and 99.34% respectively. This approach made it possible to recognise individuals from gait, relying on approximately two seconds of recording. Furthermore, these results indicate that our approach nearly always correctly identifies our participants.

**Figure 5.**
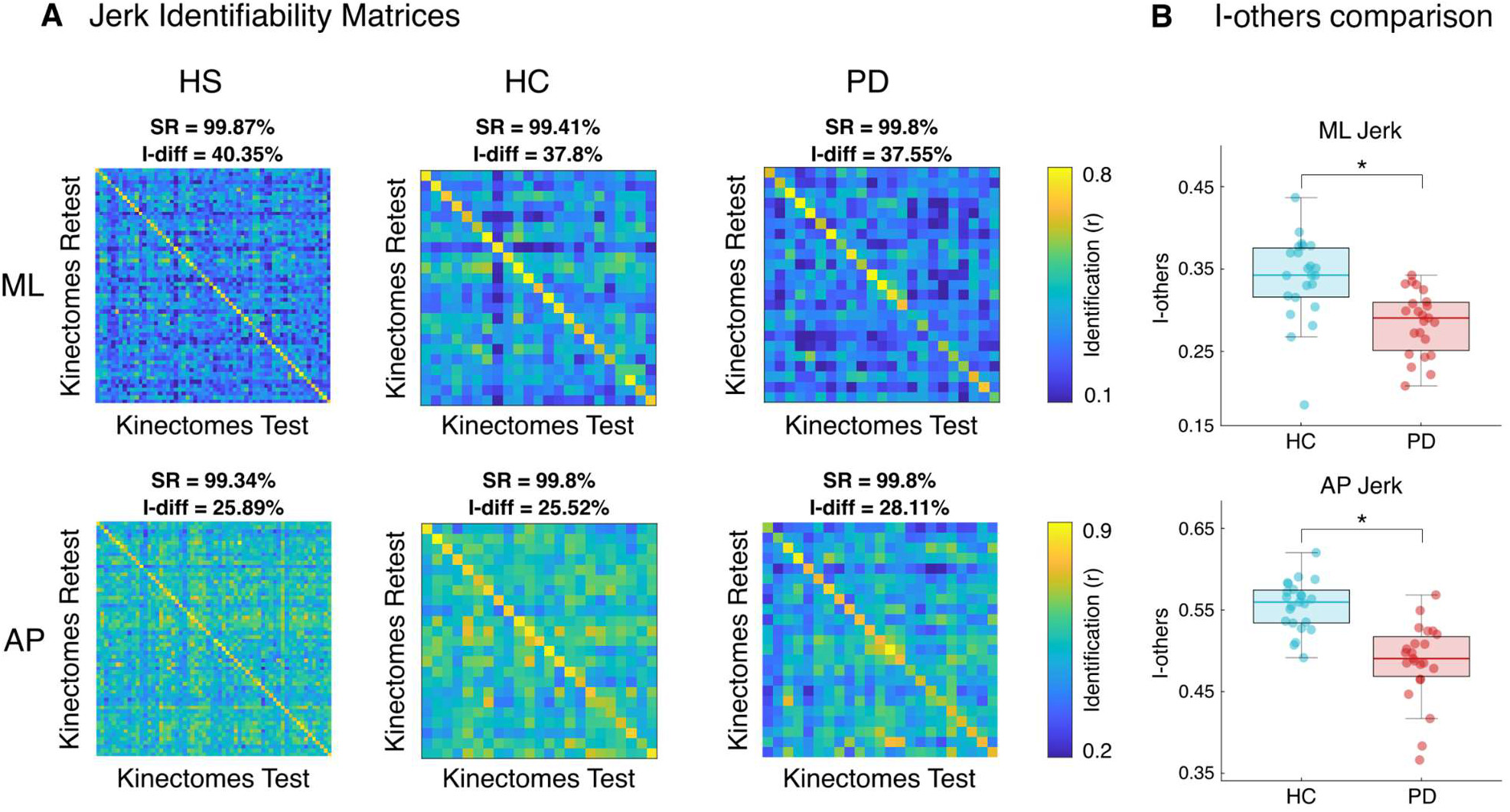
Motion fingerprinting - Identifiability based on kinectomes. **(A)** Identifiability matrices of healthy subjects (HS), healthy controls (HC) and Parkinson’s disease (PD) patients, based on jerk kinectomes in mediolateral (ML) and anteroposterior (AP) axes. Highest values within the main diagonal convey successful subject identification. **(B)** Box plot for the comparison of the I-others between HC and PD. Low I-others values indicate less within-group similarity among the subjects. The box represents data from the 25 to the 75th percentiles; the horizontal line shows the median; error lines indicate the 10th and 90th percentiles. ‘*’ represents significant Bonferroni-corrected p-values.

Comparing PD and HC groups (Bonferroni cut-off p < 0.0042), no difference was found in the I-self and I-diff. However, with respect to the I-others scores, the HC group showed higher values compared to PD patients (Fig. 5B) in AP acceleration (p < 0.0001), ML jerk (p < 0.0001) and AP jerk (p < 0.0001). Hence, the PD patients displayed more heterogeneous motor patterns with respect to the HC groups. However, concerning the identifiability matrices, both groups expressed I-diff and SR values similar to the HS group (PD ML Jerk: I-diff = 37.55%; SR = 99.8%. PD AP Jerk: I-diff = 28.11%; SR = 99.8%. HC ML Jerk: I-diff = 37.8%; SR = 99.41%. PD AP Jerk: I-diff = 25.52%; SR = 99.8%). This result highlighted that an almost complete recognition is possible for PD patients as well as for the controls.

Furthermore, we asked how this fingerprint analysis would perform when considering gait recordings performed on different days? To do this, for 11 participants we also used a second gait acquisition at a temporal distance (from the first acquisition) ranging from 19 to 164 days. Despite a slight decrease, the I-self, I-others, and I-diff scores did not show any statistical difference between the same-time gait recordings and the long-time gait recording. Moreover, even calculating the fingerprint between kinectomes related to the same subjects in two different time points, the ML Jerk showed an I-diff value equal to 28.52% and a SR equal to 98.18%; the AP Jerk identifiability displayed an I-diff equivalent to 18.85% and a SR equal to 96.36% (see Fig.S3). The analysis confirmed that gait fingerprinting is a reliable and stable feature that is maintained over time.

### Body network topology

We performed a topological analysis to examine the human movement network characteristics in a clinical setting. To this end, based on the kinectome, we computed the weighted degree (the weighted sum of all edges incident upon a given node) of each bone marker. The comparison between PD and HC (Bonferroni cut-off p < 0.0006) networks showed a significant difference in the degree of both right (p = 0.0005) and left (p = 0.0005) forearms in AP acceleration, with the HC group showing higher values compared to PD patients (Fig. 6A). Furthermore, we correlated the degree values with the Trunk displacement index (TDI), a gait stability index that was validated in PD in a previous study (*29*). Figure 6B shows the correlations between the TDI and the ML acceleration degree of C7 vertebrae (p = 0.0003; r = 0.69), T10 vertebrae (p < 0.0001; r = 0.89), left shoulder (p = 0.0003; r = 0.69), right shoulder (p = 0.0003; r = 0.69) and right elbow (p = 0.0005; r = 0.67). UPDRS scores showed a significant trend in the correlation with the ML acceleration degree of T10 vertebrae (p = 0.0007; r = 0.65). Analysing the topology of the human kinematic allowed us to highlight the main differences between PD patients and matched controls. Indeed, considering each bone marker as a node of a network, it was possible to obtain information on specific body parts, without losing the biomechanical meaning that such an element represents for the whole-body.

**Figure 6.**
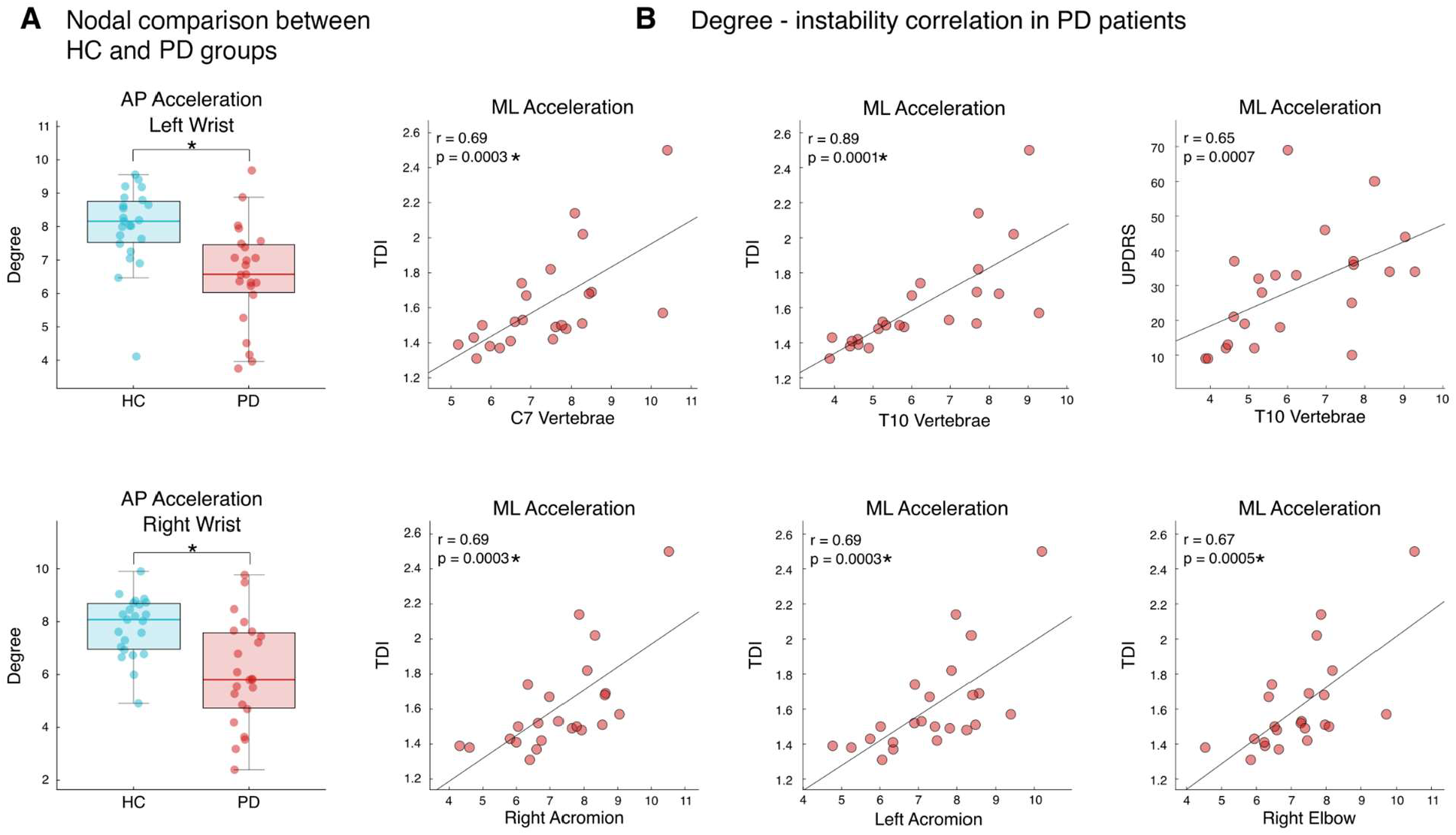
Body network examination. **(A)** Results from the nodal comparison between healthy controls (HC) and individuals affected by Parkinson’s disease (PD). The nodal degree of both wrists is lower in patients compared to controls. **(B)** Correlations between motor indices (trunk displacement index (TDI), a biomechanical index of stability, and Unified Parkinson’s Disease Rating Scale part III (UPDRS), a clinical evaluation score) and bone markers degree values. Note that most of the anatomical elements which showed a significant correlation with the instability belong to the axial component of the human body, which is commonly impaired in Parkinson’s disease. ‘*’ represents significant Bonferroni-corrected p-values.

## Discussion

In this paper we propose a novel approach to analyse human movement, based on building and investigating the mathematical structure conveying all the pairwise interactions occurring between different body segments during gait. To this end, we evaluated the covariance matrix of the accelerations of all the segments, here referred to as “kinectome”. Firstly, we showed that the kinectome provides a thorough description of the gait that distinguishes population-specific features (Fig. 2-3). Moreover, the kinectome captures symmetries in the modularity of the kinectome that are disrupted in PD patients (Fig. 4). Furthermore, through the kinectome analysis, it is possible to identify subjects based on their gait movement data obtained from two-second-long recordings (Fig. 5). Finally, the topological analysis allows us to explore the role of individual biomechanical elements of the human kinematic network. These findings confirm the utility of the kinectome in conveying the complex dynamics arising during human movement.

The presence of groups of functionally related body parts emerged naturally from the analysis of the kinectome, showing that the covariance of accelerations conveys biomechanically meaningful information. Furthermore, these patterns are specific to the clinical picture, allowing the distinction between healthy individuals and patients with PD. Notably, the best distinction between the groups was obtained using the variance of the acceleration (Fig. 3), whereby patients have much more variability as compared to the control group, and especially so with regard to the upper body, along the anteroposterior axis. We speculate that in the healthy conditions patterns of movements are optimally constrained while the neurological impairment in PD compromises the motor control, with more variable, dysregulated gait patterns appearing.

From the biomechanical perspective, the clustering analysis of the kinectome reveals the large-scale functional organization of the body segments (Fig. 4). The mediolateral allegiance matrix (that takes into account information on both the acceleration and the jerk consensus matrices – see methods for further details) of all healthy subjects (i.e., both the healthy group and the control group) showed strong coupling between the upper arms and the upper trunk, while the head, the pelvis and the forearms formed a separate functional module. Finally, the legs and the feet formed a further module. In particular, the sway of the trunk is strictly regulated and shows, in the healthy, a small range of motion during gait. This is a computationally parsimonious way to keep balance while the whole-body oscillates during gait, as well as a way to stabilize the head (*31, 32*). Furthermore, modularity analysis of the kinectome highlights the consensual accelerations among the head, the forearms, and the pelvis. The forearm might belong to this functional group since their mediolateral swing counterbalances the displacement of the centre of mass occurring during gait, thereby contributing to the maintenance of the vertical alignment (*33*). Moreover, the pelvis smoothens the movement of the COM (*34*), preventing a sharp drop toward the side of the swinging leg (*35*). Finally, the legs and the feet constitute a homogeneous community characterized by remarkable stability across the mediolateral axis. The modularity structure that we described emerges from the kinectome of the healthy individuals (including both HS and HC groups), suggesting that it might be capturing the conserved features of the healthy gait. In fact, the modularity is invariably altered in PD patients. With regard to the PD allegiance ML matrix, the modules that were observed in the healthy are altered in PD, with the body movements appearing as more fragmented. Indeed, differently from the healthy model, in PD the left and right upper arms belong to two different modules, as well as the pelvis (where upper and lower pelvis do not belong to the same functional module). These alterations may be due to the postural abnormalities (*25*) and/or to the asymmetries (*36*) induced by the pathology.

With regard to the AP axis, the allegiance matrix of the healthy groups showed three communities: the first one composed by the head, the upper trunk, and the pelvis; the second one encompassing the left leg and foot, and the right upper arm and forearm; the third community hinging on the right leg and foot, and the left upper arm and forearm. In this case, the modularity analysis separated the passenger and the locomotor units (*37*). The former is composed of the head, the trunk (including the pelvis) and the arms, the latter involves the lower limbs. However, our analysis grouped together the accelerations of the arms and the legs, characterizing two separate communities encompassing contralateral arms and legs. This separation is coherent with the fact that arms oscillate in anti-phase with respect to the contralateral legs (*38*). Interestingly, this linear pattern fails in PD, especially with respect to the trunk. In fact, the first community is composed of the head and the upper trunk, while the lower trunk and the pelvis belong to a separate community together with the right forearm. The two remaining communities constitute the anti-phased oscillations between contralateral arms and legs, as observed in the healthy groups. Once again, we can relate these disrupted patterns to the motor characteristics of parkinsonian patients. On the one hand, the asymmetry (*36*) may have caused the dysregulation of the acceleration of the right arm with respect to the healthy pattern. On the other hand, the axial rigidity, a semiological feature of the disease (*39*), does not allow the trunk to effectively relay multiple body parts. Hence, different subsections of the trunk remain entrained to more peripheral anatomical parts. In turn, this is captured by the fact that the trunk is split in different communities in patients, instead of being a coherent functional unit as seen in the HC group. Hence, the kinectomes allow us to identify features of gait that are shared by all healthy subjects and that are lost in PD patients.

However, the kinectomes can be exploited further, as to identify subject-specific gait features, thus defining a “fingerprint” of the human gait. Our analysis demonstrated that the correlations of the jerk (change in acceleration) of pairs of body segments form a unique pattern for each individual. In fact, using (approximately) two-seconds long acquisitions as test and retest sessions, we were able to identify subjects with an accuracy rate of 99% (Fig. 5A). PD patients also exhibited an identifiability rate similar to that of the controls. However, the similarity within the PD group (as measured by I-others) was lower than that within the control group (Fig. 5B). In other words, controls are more similar to each other than PD patients are. Again, one might speculate that a correct motor control imposes strict constraints to the kinectome structure, which in turn produces more similar motion patterns. In a pathological condition such as PD, such control mechanisms would fail, the constraints on the gait pattern would be loosened and, hence, the patterns would become less similar to each other.

The same experiment was reproduced in a smaller sample of healthy individuals where we identified subjects based on acquisitions performed at distant timepoints, up to five months apart. In this last case, despite a small decrease, the identification was successful even for individuals whose retest session was recorded after five months. This analysis was also based on two-seconds of walking. Although this analysis is not addressed to forensic science, this outcome can offer new insight to this field and further investigations may enhance and refine this approach. In fact, while gait identification drew considerable attention, to date the results are not robust enough for forensic applications (*40*).

Within the clinical framework, the conceptualisation of the human gait as a network encompassing the whole body, allows to quantify the contribution of single segments with respect to the large-scale patterns of movements. In fact, when representing markers as nodes and correlations of accelerations as edges, the degree of the given node will convey its overall coordination with respect to the movement of the whole body. Using this approach, the forearms emerged as less coherent in PD patients during movement across the AP axis (Fig. 6A). This result might be capturing the reduced arm swing typical of PD (*41*). The reduction of the oscillations of the upper limbs during gait has been investigated as a possible early sign of the disease (*42*–*44*). Further longitudinal studies, focusing on topological analyses of the kinectome in patients with early PD, may explore the potential of this approach in diagnostics and assessment of therapeutic responsiveness. Furthermore, in PD patients, we found several correlations between instability (measured through TDI) and the degree (based on the kinectome on the mediolateral axis) of different elements of the passenger unit, with a main involvement of the upper trunk (Fig. 6B). These correlations show that the more PD patients were unstable, the more the accelerations of the upper trunk were coherent with those of the other body segments. Axial rigidity and postural abnormalities are typical features of PD that might reflect themselves into such “hyperconnected” patterns (*45*–*47*). Intuitively, a more rigid upper trunk would require increased oscillation of the upper body during gait to maintain posture. Furthermore, the ML acceleration degree of the T10 vertebrae showed a significant trend in the correlation with the UPDRS score, once again highlighting the importance of the trunk rigidity in PD patients.

It should be stressed that, while identifiability analysis showed interesting results, the testing at distant time points was performed on eleven subjects only. Hence, this part of the results is to be considered exploitative in nature, and requires replication in larger populations. Furthermore, this is the first time that the kinectome is investigated. Hence, its ability to convey the individual clinical condition needs to be tested in samples including more PD patients, as well as both in different basal ganglia diseases and in nervous pathologies involving other systems (e.g., cerebellum, peripheral nerve or muscle). Methodologically, further analysis should be performed to explore the required number of bone markers for an optimal spatial resolution of the kinectome. Finally, in this work we only consider pairwise interactions, future work should also consider higher-order interactions (*48*).

In conclusion, we introduced a new mathematical tool, the kinectome, to convey general gait patterns in humans, subject-specific motor features and, finally, features that are disease-specific. This approach revealed several potentially useful pieces of information. Movement fingerprinting may be further exploited for security purposes as well as to longitudinally monitor individual gait features. Using topological metrics we were able to localize some of the main changes occurring in PD patients. Further studies are necessary to investigate the clinical potential of the kinectome. Finally, the use of the kinectome analysis may be of help in both sport training and physiotherapy. We hope that our work will contribute to the development of mathematical approaches to describe human movements within the “kinectomics” framework, and toward the representation of human movement as a complex integrated system.

## Materials and Methods

### Participants

Sixty healthy subjects including 38 males and 22 females were recruited (mean age 58.7 ± 12.7 years). Exclusion criteria were the following: (a) Mini-Mental State Examination (MMSE) < 24 (*49*); (b) Frontal Assessment Battery (FAB) < 12 (*50*); (c) Beck Depression Inventory II (BDI-II) > 13 (*51*); neurological or psychiatric disorders; (e) intake of psychoactive drugs; (f) physical or medical conditions causing motor impairment.

To test the validity of our methods in a clinical setting, we used the data of twenty-three patients (mean age 65.3 ± 11.6) affected by Parkinson’s disease and twenty-three healthy controls, matched for age, sex and education. The subjects included in this study are partially overlapping with those included in Troisi Lopez et al. (*29*). Parkinsonians were tested in off-medicament state. Inclusion criteria were: (a) Hoehn and Yahr (H&Y) score ≤ 3 while off-medicament (*52*); (b) disease duration < 10 years; (c) antiparkinsonian treatment at a stable dosage. All participants signed an informed consent in accordance with the declaration of Helsinki. The study was approved by the “Azienda Ospedaliera di Rilievo Nazionale A. Cardarelli’” Ethic Committee (protocol number: 00019628).

### Stereophotogrammetric acquisition

The acquisitions were carried out in the Motion Analysis Laboratory of the University of Naples Parthenope. Gait data were recorded through a stereophotogrammetric system for motion analysis composed of eight infrared cameras (ProReflex Unit—Qualisys Inc., Gothenburg, Sweden), capturing (at 120 frame per second) the light reflected by 21 passive markers positioned on the naked skin of the participants. The markers were placed in correspondence of bone landmarks, based on a modified version of the Davis protocol (*53*). We asked the participants to walk in a straight path choosing their preferred walking speed. For each participant, two gait acquisitions were performed, each of which included one complete left and right gait cycle. A complete gait cycle is defined as starting with the heel touching the ground, and finishing with the next contact with the ground of the same heel. Eleven healthy participants underwent a second recording several days after the first one (ranging from 16 to 164 days), to test the reliability of the gait fingerprinting. Through the Qualisys Track Manager software we obtained the three-dimensional position of each bone marker during the gait cycle. Hence, we could calculate the time series for acceleration and jerk (the first derivative of acceleration with respect to time) of each bone marker.

### Introducing the kinectome

To obtain the kinectome, we computed Pearson’s correlation coefficients between the bone markers’ time series (see also Fig. 1A - 1B), obtaining a covariance matrix. Specifically, we computed a kinectome for the bone markers’ acceleration and jerk separately as well as for two different movement directions (i.e., mediolateral and anteroposterior). Firstly, we explored the kinectome heterogeneity within and between groups (PD patients and controls), by comparing mean and standard deviations of the kinectomes. We then characterized the kinectomes utilizing graph-theoretical analyses, as detailed in the next sections.

### Modularity analysis

Modularity is a measure of the strength of division of a network into modules or communities. Networks with high modularity have dense connections between the nodes within modules but sparse connections between nodes in different modules. We assessed the community structure (i.e., partition) of each group-averaged kinectome (anteroposterior and mediolateral, separately), in both healthy and PD patients, by using the Louvain method (*54*) for identifying communities in large networks. In order to improve the stability of the community detection procedure, we performed consensus clustering (*55*) out of a set of 100 partitions obtained with the Louvain method. The consensus clustering technique performs a search for a consensus partition, i.e., the partition that is most similar, on average, to all the input partitions (Fig. 1C). While the similarity can be measured in several ways, in this work we chose the probability of co-occurrence of the nodes within a specific community (*55*), i.e. the allegiance matrix (*30*). Indeed, we computed the modularity of the allegiance matrix for each axial direction of movement. To do so, for each axial direction, a consensus matrix was built including acceleration and jerk at once, as to condense the information contained in both quantities. The aim was to identify, in a data-driven fashion, functional dynamical clusters within healthy and diseased kinectomes during gait.

### Fingerprint analysis

Can we identify individuals based solely on their motion patterns, i.e., their kinectomes? To address this question, we took inspiration from previous studies on fingerprint in human functional brain connectomes extracted from fMRI and MEG data (*26, 27*). In a recent work (*26*), the authors defined a mathematical object known as identifiability matrix, which encodes the information about the self-similarity (I-self, main diagonal elements) of each subject with herself/himself, across the test/retest sessions, and the similarity of each subject with the others. In order to build an identifiability matrix based on kinectomes, we first considered two gait cycle registration for each individual, called test and retest respectively. We then obtained the identifiability matrix through Pearson’s correlation between test and retest of our subjects (Fig. 1D). The main diagonal of this matrix contains the similarity between two separate acquisitions of the same subject (self-similarity or I-self); the off-diagonal elements contain the similarity between each subject with the test or retest acquisition with respect to the other subjects (I-others). Furthermore, the difference between I-self and I-others, also known as differential identifiability (I-diff), provides a robust score of the overall fingerprinting assessment of a dataset. Finally, we estimated the success rate as the percentage of times in which an in-diagonal element coefficient was higher than the out-diagonal elements coefficients belonging to the row and column of the in-diagonal element taken into consideration.

### Topological analysis of the kinectome

We represented the body as a network, where body parts are nodes and their correlations form the edges, obtaining a weighted undirected graph (Fig. 1E). For each graph, we estimated the weighted degree, a centrality parameter (*15, 56*). The degree was calculated as the sum of the absolute value of the edge weights for each node (*57*).

### Statistics

Statistical and data analysis were carried out in MATLAB 2020a. Significance of the between groups (PD and HC) differences in the kinectomes standard deviation, fingerprint values (I-self, I-other, I-diff), and topological parameter (degree) were assessed through permutation testing, by randomly shuffling group labels 10000 times. At each permutation, the absolute value of the difference was computed, obtaining a distribution of the differences that are to be expected by chance alone (*58*). This distribution was compared to the observed differences to retrieve a statistical significance. Correlation analysis between nodal degree and motor scores was performed through the Spearman correlation test. The significance threshold was set at *p* < 0.05, and was Bonferroni corrected to adjust for multiple comparisons in each analysis.

## Notes

### Competing Interest Statement

The authors have declared no competing interest.

