## Supplementary materials for "The kinectome: a comprehensive kinematic map of human motion in health and disease"

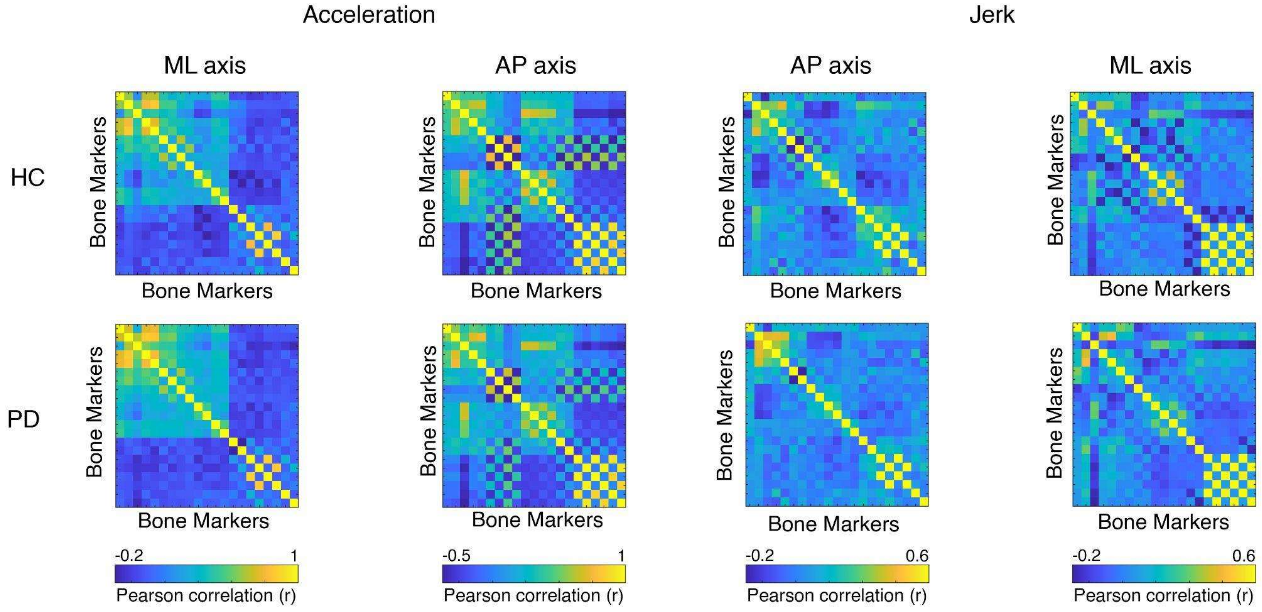


**Figure S1. Group average kinectomes in patients and controls**

Averaged kinectomes for the Parkinson's disease group (PD) and the healthy control group (HC). Acceleration and jerk kinectomes are shown, for both mediolateral (ML) and anteroposterior (AP) axes. Despite slight divergences, the kinectomes appear similar between the two groups and no statistical difference was found.

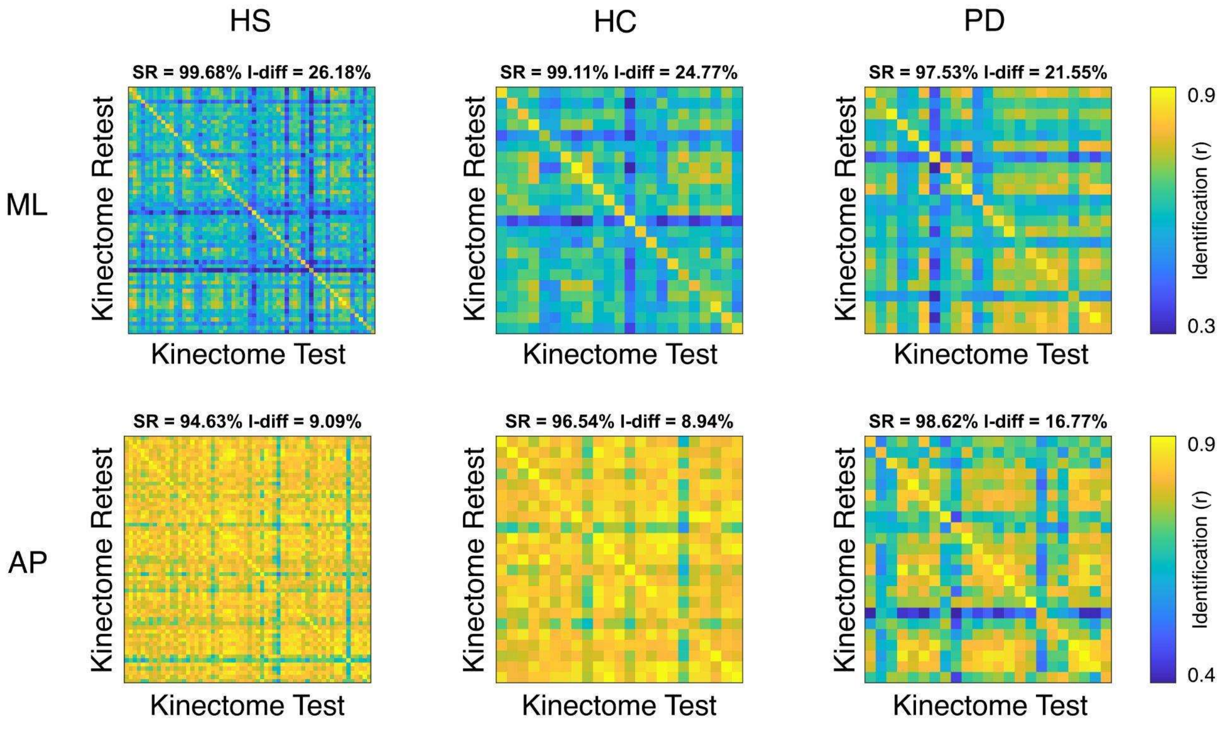


**Figure S2. Fingerprint analysis in acceleration-based kinectomes**

Identifiability matrices for the acceleration-based kinectomes of healthy subjects (HS), healthy controls (HC) and Parkinson’s disease (PD) groups. Similar to identifiability matrices based on jerk kinectomes, the mediolateral axis performs better in identifying subjects.

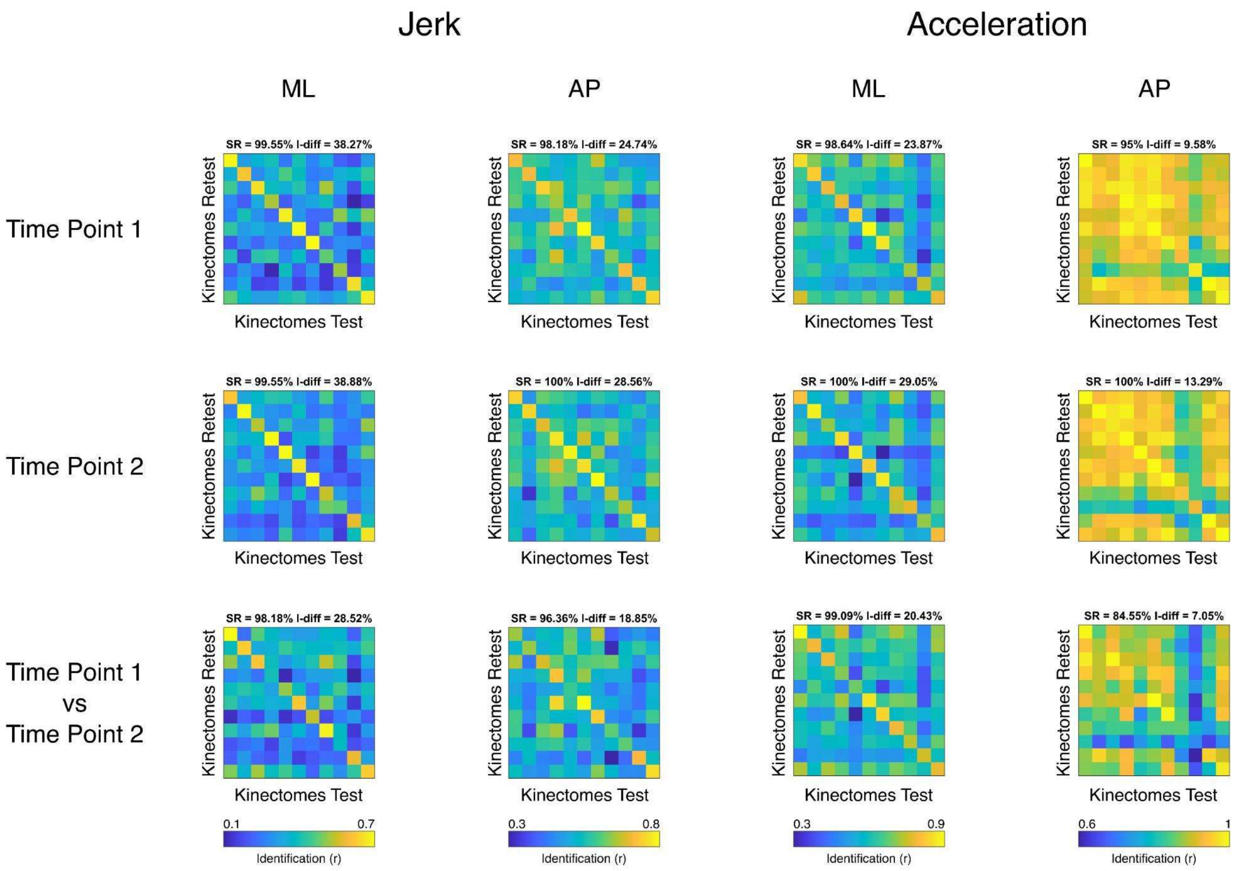


**Figure S3. Longitudinal identification**

Identifiability matrices for subject recognition, based on gait acquisition recorded in different time points. Results for jerk and acceleration are shown in mediolateral (ML) and anteroposterior (AP) axes. The first row shows the identifiability matrices accounting for test-retest kinectomes of gait cycles acquired within the same day (Time Point 1). The second row too displays the identifiability matrices referring to acquisition occurred within the same day but at a temporal distance (with respect to the first acquisition) ranging from 19 to 164 days. Finally, the third row shows the identifiability matrices obtained comparing kinectomes of the same subjects recorded at different time points. Note that, despite a slight decrease, both success rate (SR) and I-diff values are not significantly different when comparing the longitudinal identification (row three) with the within same day identification (first and second rows).
